# Nest-building using place cells for spatial navigation in an artificial neural network

**DOI:** 10.1101/2023.05.30.542884

**Authors:** Thomas E. Portegys

## Abstract

An animal behavior problem is presented in the form of a nest-building task that involves two cooperating birds, a male and female. The female builds a nest into which she lays an egg. The male’s job is to forage in a forest for food for both himself and the female. In addition, the male must fetch stones from a nearby desert for the female to use as nesting material. The task is completed when the nest is built and an egg is laid in it. A goal-seeking neural network and a recurrent neural network were trained and tested with little success. The goal-seeking network was then enhanced with “place cells”, allowing the birds to spatially navigate the world, building the nest while keeping themselves fed. Place cells are neurons in the hippocampus that map space.

## Introduction

A task is presented to simulate two birds, a male and a female, that cooperate in navigation, foraging, communication, and nest-building activities. These activities are commonly performed in many animal species to ensure survival and reproduction. The female builds a nest into which she lays an egg, completing the task. The male’s job is to forage in a forest for food for both himself and the female. In addition, the male must fetch stones from a nearby desert for the female to use as nesting material.

The nest-building task was recently proposed as a game providing an artificial animal intelligence challenge to evaluate machine learning systems (Portegys, 2022a). While the task itself is independent of how it is tackled, here artificial neural networks (ANNs) are chosen, as ANNs are capable of generalized learning, and are intended to perform functions of the biological neural networks of animals.

The task was originally introduced in 2001 (Portegys, 2001), a solution for which was obtained using Mona, a goal-seeking ANN that at the time employed domain-specific macro-responses, such as “Go to mate”. The network was also manually coded instead of being learned.

In planning to attack the problem again, this time as a learning task with spatial enhancements (Portegys, 2022b) that would obviate the need for domain-specific responses, place cells seemed to be a good choice. Place cells are neurons in the hippocampus that map space (Moser et al., 2015; Robinson et al., 2020; Xu et al., 2019) allowing an animal to navigate its environment effectively.

There is a significant body of work on using place cell inspired neural functionality in ANNs, much of it involved with robotic navigation (Milford and Wyeth, 2010; Zhou et al., 2017). These systems are aimed at solving specific tasks with models that mimic biological place cells. They are not intended to be general-purpose ANNs, such as Mona, which are designed to learn arbitrary domain-independent tasks. General-purpose ANNs borrow functionality from brains, such as neural connection updating, but are not intended to be models of biological brains. Here place cells are incorporated into an ANN to allow it to effectively operate in spatial environments. To my knowledge this is a novel development.

As a comparison, an LSTM (Long short-term memory) recurrent neural network (RNN) (Hochreiter and Schmidhuber, 1997) was also trained on the task, without spatial enhancement.

Historically, AI has mostly focused on human-like intelligence, for which there are now numerous success stories: games, self-driving cars, stock market forecasting, medical diagnostics, language translation, image recognition, etc. The impact of ChatGPT (OpenAI, 2023) as a generative language model is a recent example. Yet the elusive goal of artificial general intelligence (AGI) seems as far off as ever, likely because these success stories lack the “general” property of AGI, operating as they do within narrow, albeit deep, domains. A language translation application, for example, does just that and nothing else.

Anthony Zador (2019) expresses this succinctly: “We cannot build a machine capable of building a nest, or stalking prey, or loading a dishwasher. In many ways, AI is far from achieving the intelligence of a dog or a mouse, or even of a spider, and it does not appear that merely scaling up current approaches will achieve these goals.”

Mona is a goal-directed ANN, designed to control an autonomous artificial organism. We can go back to Braitenberg’s vehicles (1984) as a possible starting point for the study of how simple networks are capable of controlling goal-directed behavior in automata. Dyer (1993) delineated the contrasting aims of ANNs for animal behavior vs. traditional artificial intelligence (AI) by framing the former as having biological goals: “Survivability is the overriding task”. As an example of a simulated animal brain, Coleman et al. (2005) developed an ANN-based cognitive-emotional forager that performed well in a task that required not only foraging, but also the avoidance of predators.

Perhaps the best example of artificial creatures controlled by ANNs is Yaeger’s Polyworld artificial life system (Lizier et al., 2009). Polyworld is an environment in which a population of agents search for food, mate, have offspring, and prey on each other in a two dimensional world. An individual makes decisions based on its neural network which is derived from its genome, which in turn is subject to evolution. To my knowledge, however, training an ANN to build a nest is a novel undertaking.

In addition to simulating life forms, ANNs have been used as modeling and analysis tools for animal behavior (Enquist and Ghirlanda, 2006; Wijeyakulasuriya et al., 2020).

## Description

The code and instructions can be found at: https://github.com/morphognosis/NestingBirds

### World

The world is a 21×21 two dimensional grid of cells. Each cell has a *locale*, and an *object* attribute. *Locale* describes the type of terrain: *plain, forest*, and *desert*. An *object* is an item to be found in the world: *mouse* (food), and *stone* (nest-building material). A forest exists in the upper left of the world, populated by mice, which randomly move about, providing an element of surprise for a foraging bird. A desert is found in the lower right of the world, where stones are to be found at various locations. The birds are initially located on the plain in the center of the world.

### Birds

There is a male and a female bird. The female builds the nest and the male forages for mice and stones. The nest is a stone ring in the center of the world in which the female lays her egg. The birds have four components: senses, internal state, needs, and responses. These are sex-specific to suit the different roles of the birds.

#### Male

##### Senses

*locale, mouse-proximity, stone-proximity, mate-proximity, goal, has-object, flying, female-needs-mouse, female-needs-stone*.

*Locale* pertains to the current location of the male and has a value of *plain, forest*, or *desert*.

The *proximity* sensor values are *present, left, right, forward*, or *unknown*. The *mouse-proximity* sensor senses a mouse when in the forest, the *stone-proximity* sensor senses a stone when in the desert, and the *female-proximity* sensor senses the female within the bounds of the nest.

The *goal* sensor values are *eat-mouse, mouse-for-female, stone-for-female*, and *attend-female*.

The *has-object* sensor indicates an object carried by the bird and can *be mouse, stone*, or *no-object*.

The *flying sensor* is true when the male is in flight; otherwise false.

*Female-needs-mouse* is sensed when the female expresses a corresponding response of *want-mouse* in the presence of the male. This is the only time this sense is active; when not in the presence of the female it is in the off state. A similar process exists for the *female-needs-stone* sense and *want-stone* female response. Only one of the *female-needs/wants* is sensed/expressed at a time.

##### Internal state

*food*.

Initialized to a parameterized value. When food reaches zero, the need for a mouse is surfaced as a goal. Upon eating a mouse, food is increased by a random value.

##### Needs

*mouse-need, female-mouse-need, female-stone-need, attend-female-need*.

These correspond to the goals. Upon completion of an errand to satisfy a goal, signified by returning to the female, the next goal is determined by current needs. As discussed, *mouse-need* is raised when food reaches 0. The *female-mouse-need* and *female-stone-need* are signaled to the male by the female when she desires a mouse to eat or a stone to place in the nest, respectively. If none of the above are raised, the *attend-female-need* is raised, causing the male to move to the female’s location.

##### Responses

*do-nothing*: a no-op response.

*move-forward*: move forward in the orientation direction. Movement off the grid is a no-op.

*turn-right/left*: change orientation by 90 degrees.

*eat-mouse*: eat mouse if *has-object* is a mouse. If no mouse, this is a no-op.

*get-object*: if *has-object* is empty and an object in current cell, pick up the object and set it to *has-object*.

*put-object*: if *has-object* not empty and no object at current cell, put object in cell and clear *has-object*.

*give-mouse*: if *has-object* is mouse, and female present with empty *has-object*, transfer mouse to female.

*give-stone*: if *has-object* is stone, female present with empty *has-object*, transfer stone to female.

*fly*: take flight. This activates a place motor neuron which will move to a specific location in the world. The specific motor is determined by the current mediator neuron context (see Artificial neural networks section).

*alight*: terminate flight after arriving at a location in the world determined by the active place motor neuron.

#### Female

##### Senses

*object-sensors, orientation, goal, has-object*.

*object-sensors*: the female senses the object values in its Moore (3×3) neighborhood.

*orientation*: north, south, east and west. *goal*: lay-egg, brood-egg, eat-mouse. *has-object*: identical to male.

##### Internal state

*food*.

Identical to male.

##### Needs

*lay-egg-need, brood-egg-need, mouse-need*.

The *mouse-need* need is raised by food level and sets the *eat-mouse* goal. It will cause the female to express the *want-mouse* response. While building the nest and not hungry, the *lay-egg-need* is kept raised with the associated *lay-egg* goal. The female asserts the *want-stone* response when she is located in a cell where the nest requires a stone to be placed. After a stone is placed, the female proceeds to the next location in the nest and repeats the process, continuously motivated by the *lay-egg-need* and *lay-egg* goal. When the nest is built, the female lays her egg in the center of the nest. After that *brood-egg*-*need* is kept raised and the *brood-egg* goal keeps the female brooding on the egg.

##### Responses

*do-nothing*: a no-op response.

*move-forward*: move forward in the orientation direction. Movement off the grid is a no-op.

*turn-right/left*: change orientation by 90 degrees.

*eat-mouse*: eat mouse if *has-object* is a mouse. If no mouse, this is a no-op.

*get-object*: if has-object is empty and object in current cell, pick up the object and set it to *has-object*.

*put-object*: if *has-object* and no object in current cell, put object in cell and clear *has-object*.

*want-mouse*: when the *eat-mouse* goal is sensed, the *want-mouse* response signals the male to retrieve a mouse from the forest for the female to eat.

*want-stone*: when the *lay-egg* goal is sensed, and the female is ready to place a stone in the nest, the *want-stone* response signals the male to retrieve a stone from the desert for the female to place in the nest.

*lay-egg*: when the female has completed the nest and has moved to its center, she lays the egg with this response.

#### Artificial neural networks

##### *M**ona*

Mona learns cause-and-effect chains and hierarchies of neurons that represent these relationships in the environment. A detailed description of the architecture can be found in Portegys (2001). An overview is provided here.

Three types of neurons are defined, as shown in Figure 1:

**Figure 1.**
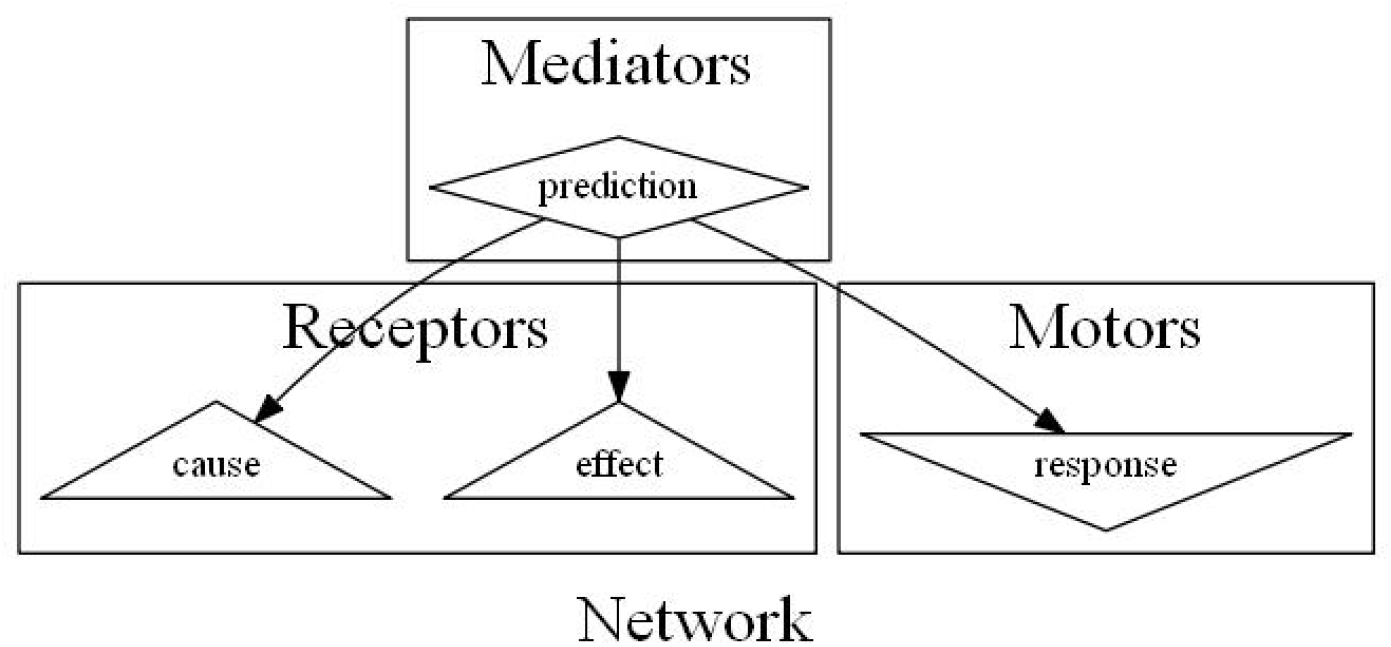
A simple Mona network

- Receptor neuron: represents a sensory event.
- Motor neuron: expresses response to the environment.
- Mediator neuron: represents a predictive relationship between neurons. The firing of its cause neuron *enables* its motor neuron to fire. If the network fires the motor neuron, the effect neuron will probabilistically fire. A mediator that mediates lower level mediators serves as a context that recursively affects the probability of causation in its components.

Mona is a goal-seeking network, falling into the category of a model-based reinforcement learning system (Moerland et al., 2023). Needs arising from internal and external sources are satisfied by the firing of associated goal neurons. For example, a need for water is satisfied by firing neurons involved with drinking water. Need drives backward through the network from a goal neuron as *motive*, following enabled pathways to fire a motor response that will navigate toward the goal.

There can be multiple needs vying for control. For example, food and thirst might both act on the network simultaneously. The winner will be a function of the strength of the need and the enablement of the network to achieve the goal.

In the nest-building task, the male learns a network consisting of chains of mediators that drives it to the forest when it needs a mouse to eat, then returns it to the female for further goal setting. It also learns mediators that orchestrate fetching mice and stones for the female. A powerful feature of Mona is that mediators are modular, that is, they can take part in multiple goal activities. For example, the mediators that fly the male to the forest and hunt for a mouse are used when both the male and female require a mouse.

The female’s need to lay an egg, in conjunction with the goal of sensing the egg in the nest, drives through a chain of mediators that orchestrate a series of nest-building actions, each of which involves expressing a want for a stone that signals the male to fetch a stone, placing the stone, and moving to the next location until the nest is entirely constructed. The female then moves to the center of the nest and lays her egg, satisfying the egg laying need. Then the need to brood the egg is raised, which keeps the female sitting on the egg.

Bird responses are trained by overriding incorrect responses with correct ones. The correct responses are incorporated into the network. During testing, responses are generated by the network.

#### Place motor neurons

To enhance Mona with place cell functionality, a place motor neuron was implemented. A biological place neuron fires when a specific location in the world is reached. Mona’s place neurons fire similarly. However, they also navigate to specific places in the world. A related capability seems to exist in the form of route encoding in rats (Grieves et al., 2016).

In Mona, a place neuron is implemented as a motor neuron that records a specific location in the world and will navigate to that location when motivated to respond. It fires when the location is reached. For example, when the male’s goal is to fetch a mouse for the female, a mediator with a cause of sensing the female’s want of a mouse, and an effect of sensing the proximity of a mouse in the forest will fire its place motor to enact a series of primitive movement responses that navigate to a prescribed location in the forest.

Place motors can be learned while exploring the world by marking significant locations. A special responses initiates the commencement of a sequence of movements that terminate at some location, at which time another special response marks the location, creating a new place motor neuron. In the nesting birds, these two special responses are mapped to the male *fly* and *alight* responses, respectively.

##### LSTM

The LSTM, introduced in 1997 (Hochreiter and Schmidhuber, 1997), is a recurrent neural network which has established itself as a workhorse for sequential pattern recognition. LSTMs address a problem with other recurrent neural networks in which corrective information vanishes as the time lag between the output and the relevant input increases, leading to the inability to train long-term state information.

In the LSTM network, the hidden units of a neural network are replaced by memory blocks, each of which contains one or more memory cells. A memory block is shown in Figure 2. The block can latch and store state information indefinitely, allowing long-term temporal computation. What information to store, and when to output that information are part of the training process.

**Figure 2.**
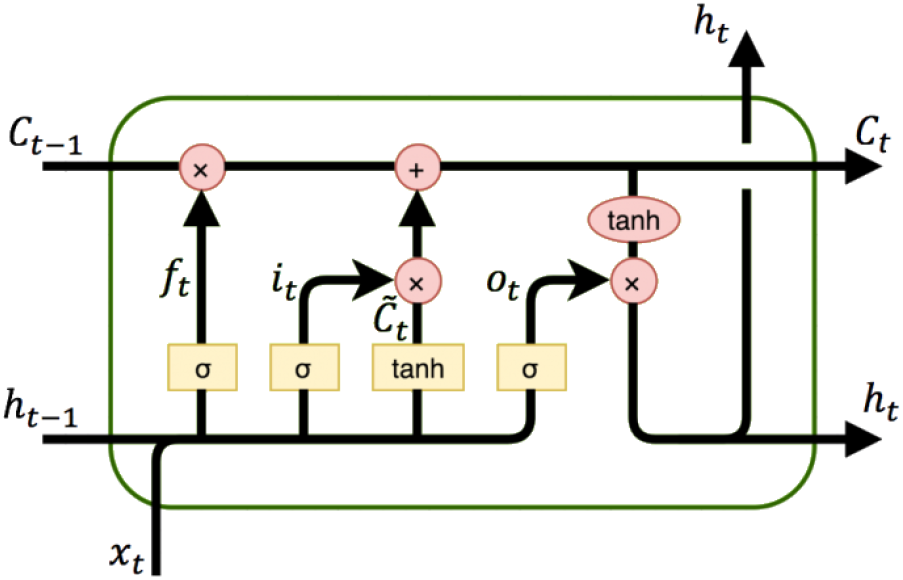
LSTM memory block.

**Figure 3.**
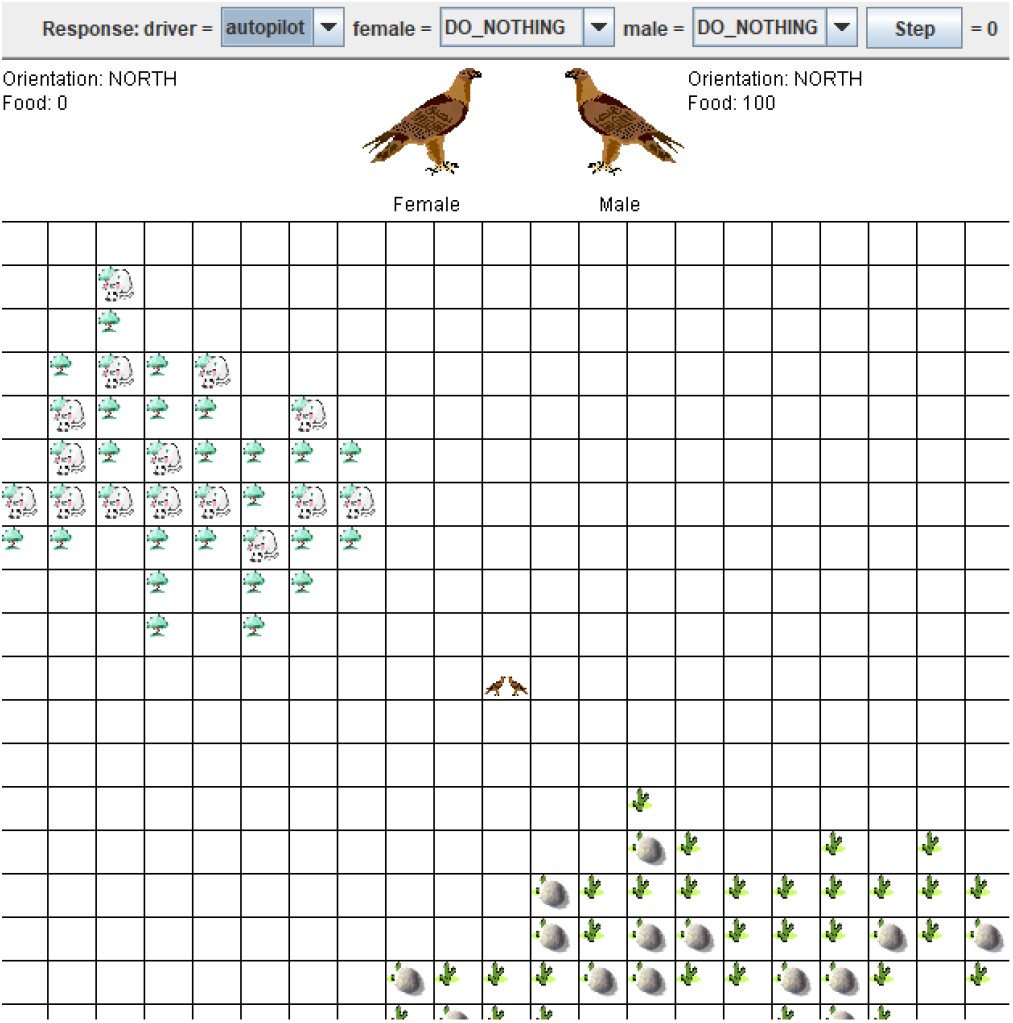
Beginning state. Female is hungry (0 food), male has maximum food. Initial response for both is “do nothing”. Both are located at center of world. Upper left is forest with mice (food). Lower right is desert with stones for nest-building.

**Figure 4.**
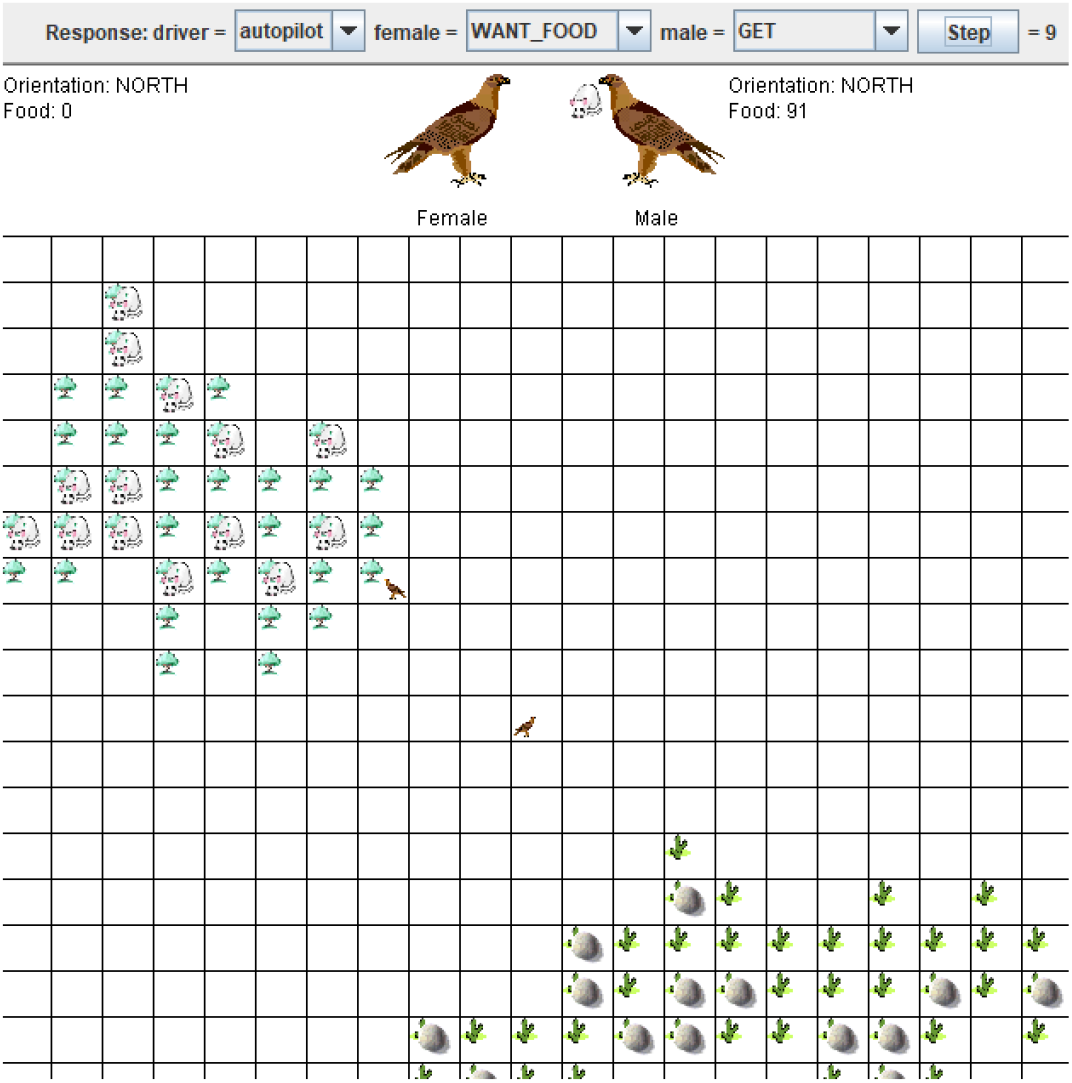
While co-located, female signals to male with “want food” response. Male flies to forest and picks up a mouse to feed to her.

**Figure 5.**
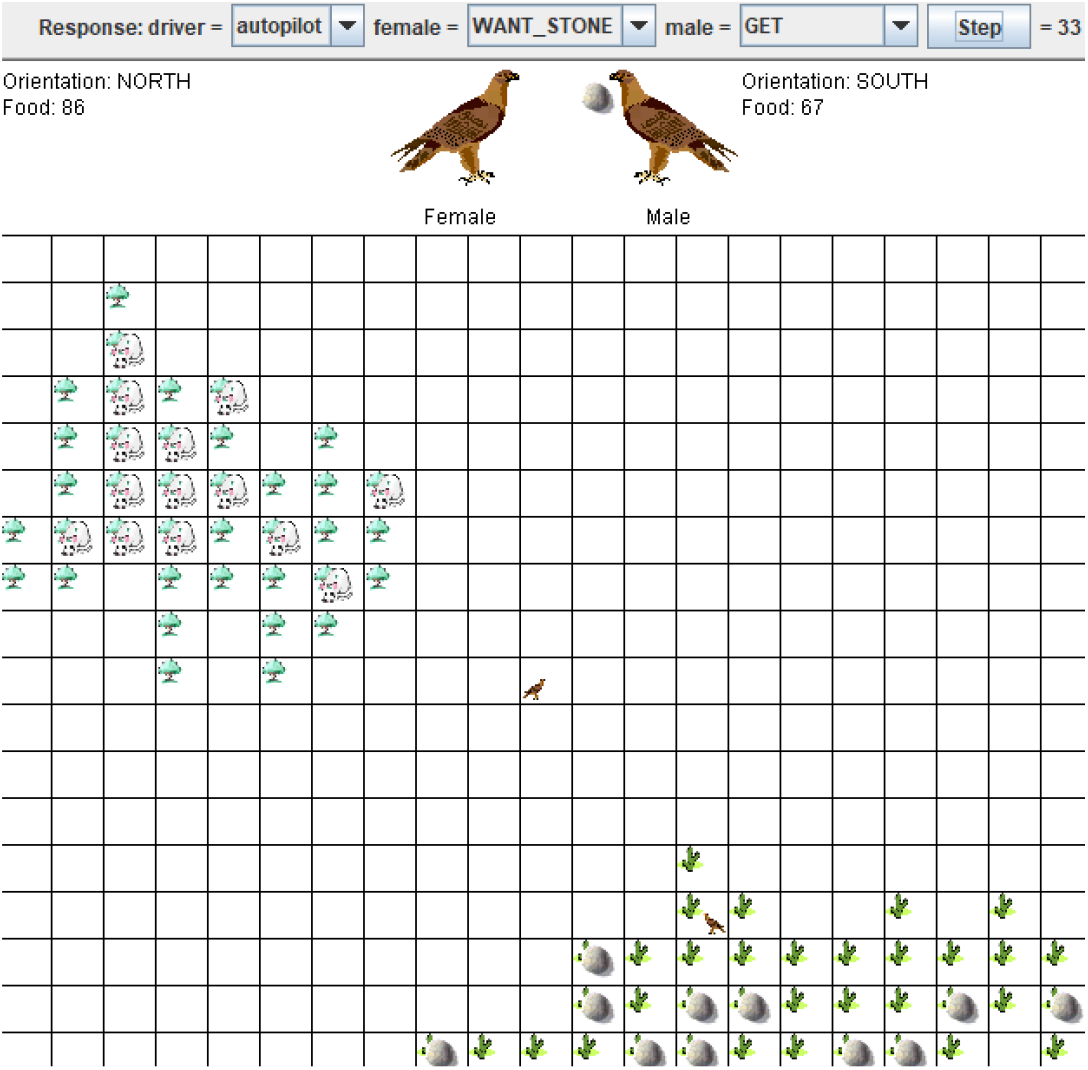
Female moves to location of first nesting stone. Male follows her. She signals to male that she wants a stone. Male flies to desert and picks up a stone.

**Figure 6.**
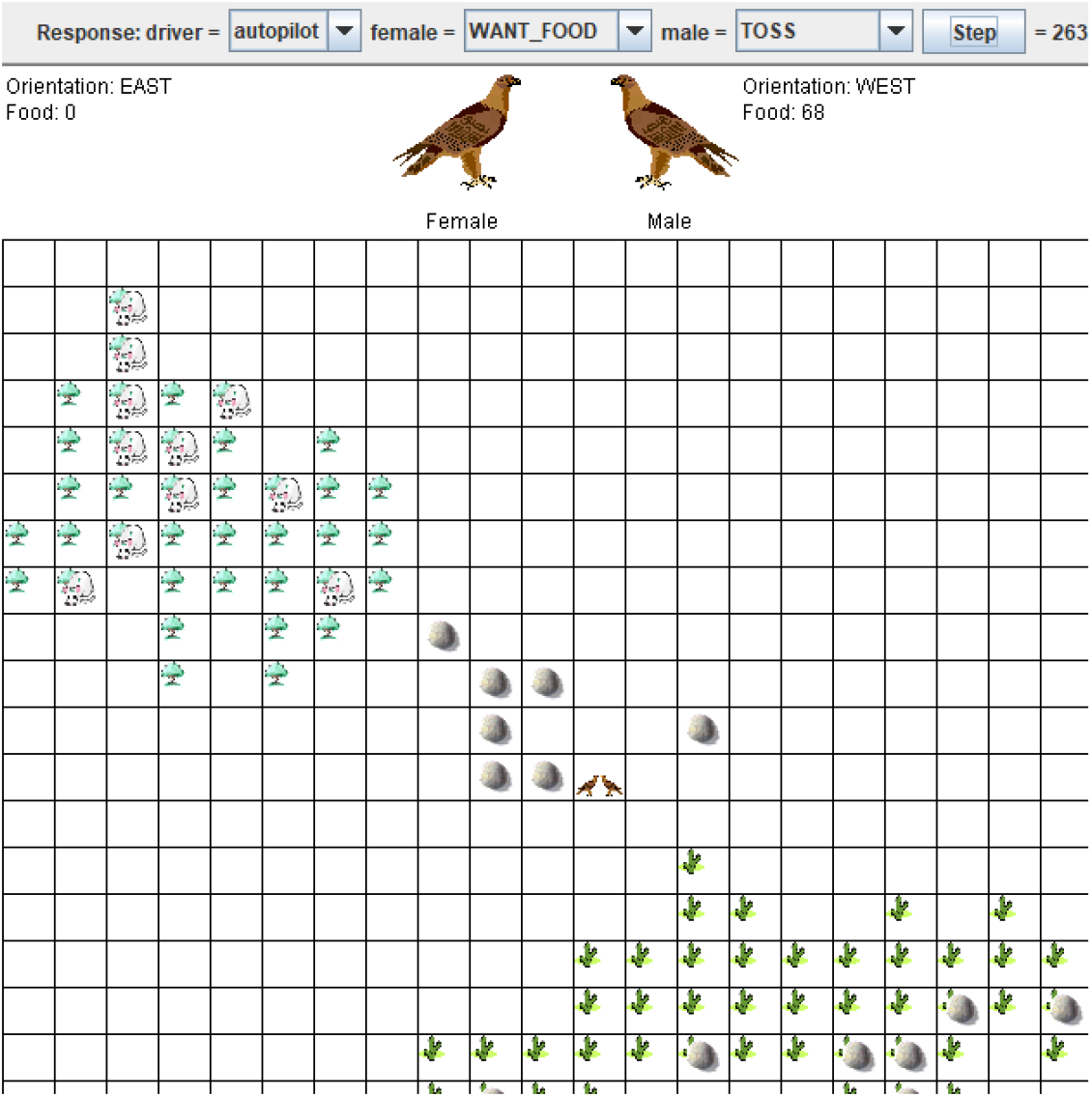
Male returns to female with stone. Discovers she is hungry. He flies to forest for mouse for her.

**Figure 7.**
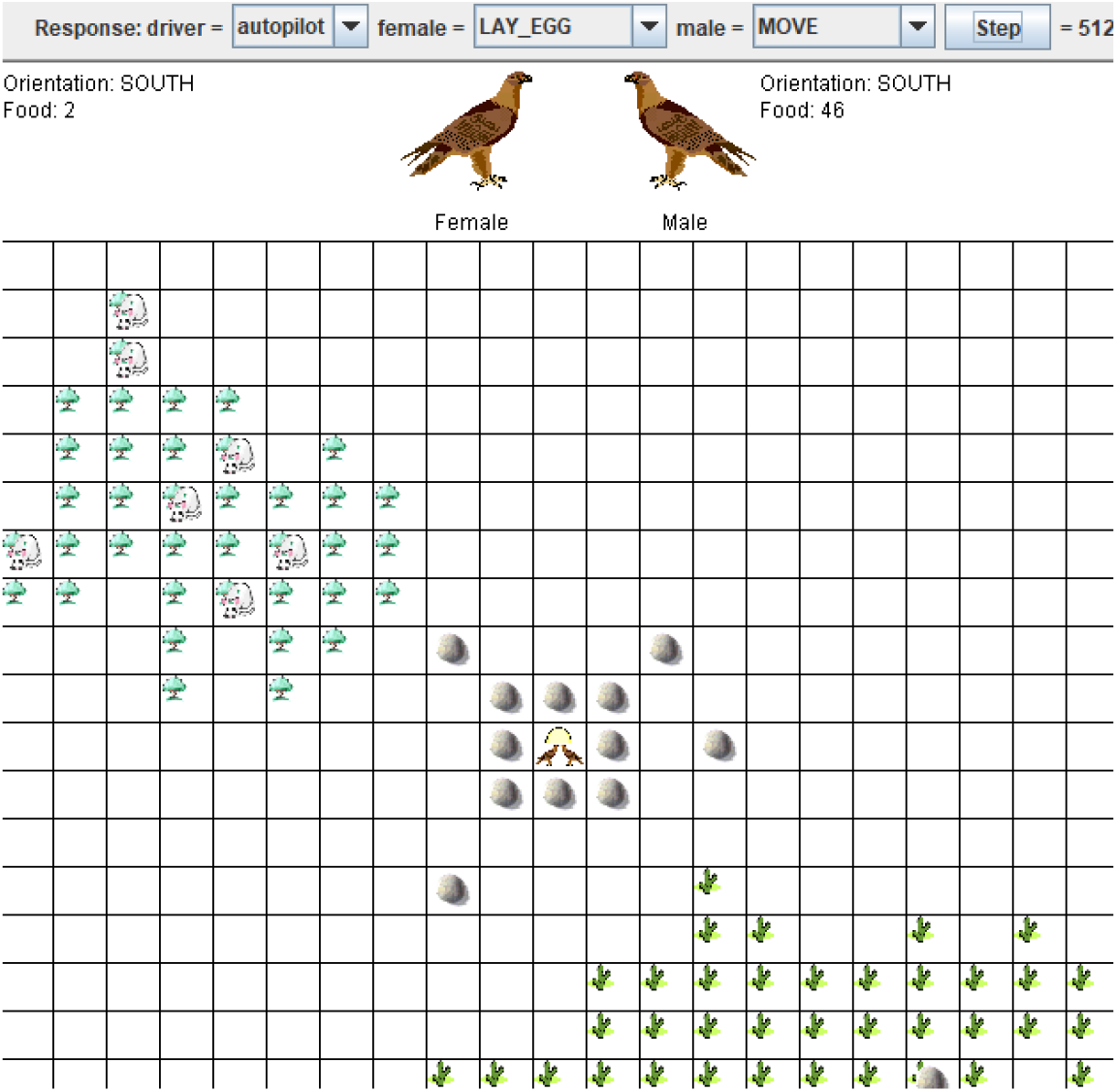
Nest completed. Egg laid.

### Scenario

This scenario is taken from the game proposal (Portegys, 2022a), which illustrates the task from a general viewpoint. It shows intermittent states of the world, from initial state to egg-laying in the completed nest. A video is available here: https://youtu.be/d13hxhltsGg

## Results

Two ANNs were trained and tested on the task under varying conditions:

1. Mona version 6.0. Maximum of 500 mediator neurons. Maximum mediator level of 0, meaning mediators mediated receptor and motor neurons exclusively; higher level mediators that mediate lower level mediators were not needed.
2. An LSTM (Long-short term memory) recurrent neural network using the Keras 2.6.0 python package. 128 neurons in a hidden layer, and a mean squared error loss function. Input and output were one-hot encoded. Training was conducted with 500 epochs.

### Number of training datasets

Performance was measured based on the number of training datasets, as shown in Figures 8 and 9. A training dataset is generated by running the nesting birds with optimal bird responses until the nest is completed while keeping the birds fed.

**Figure 8.**
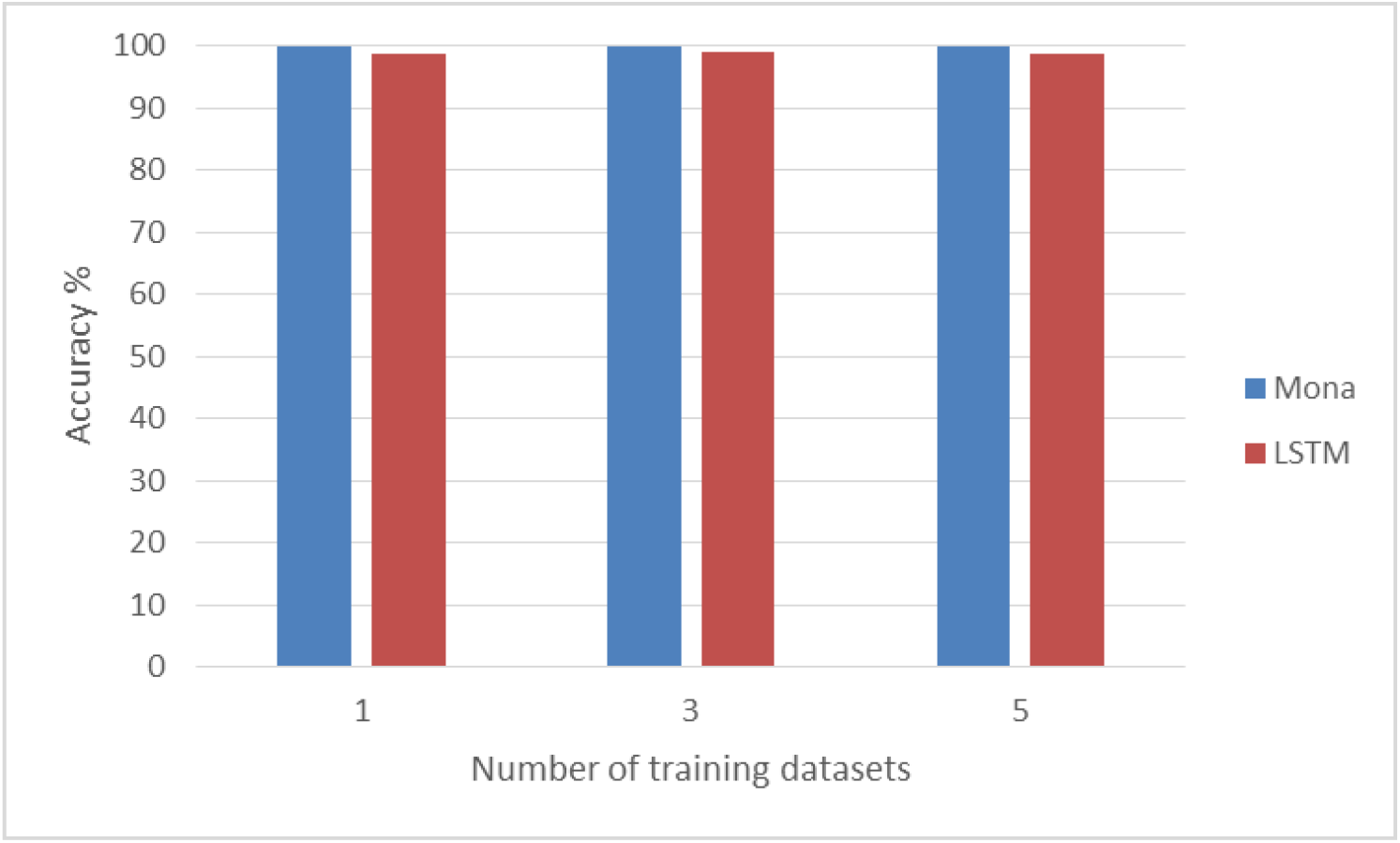
Female test performance with number of training datasets

**Figure 9.**
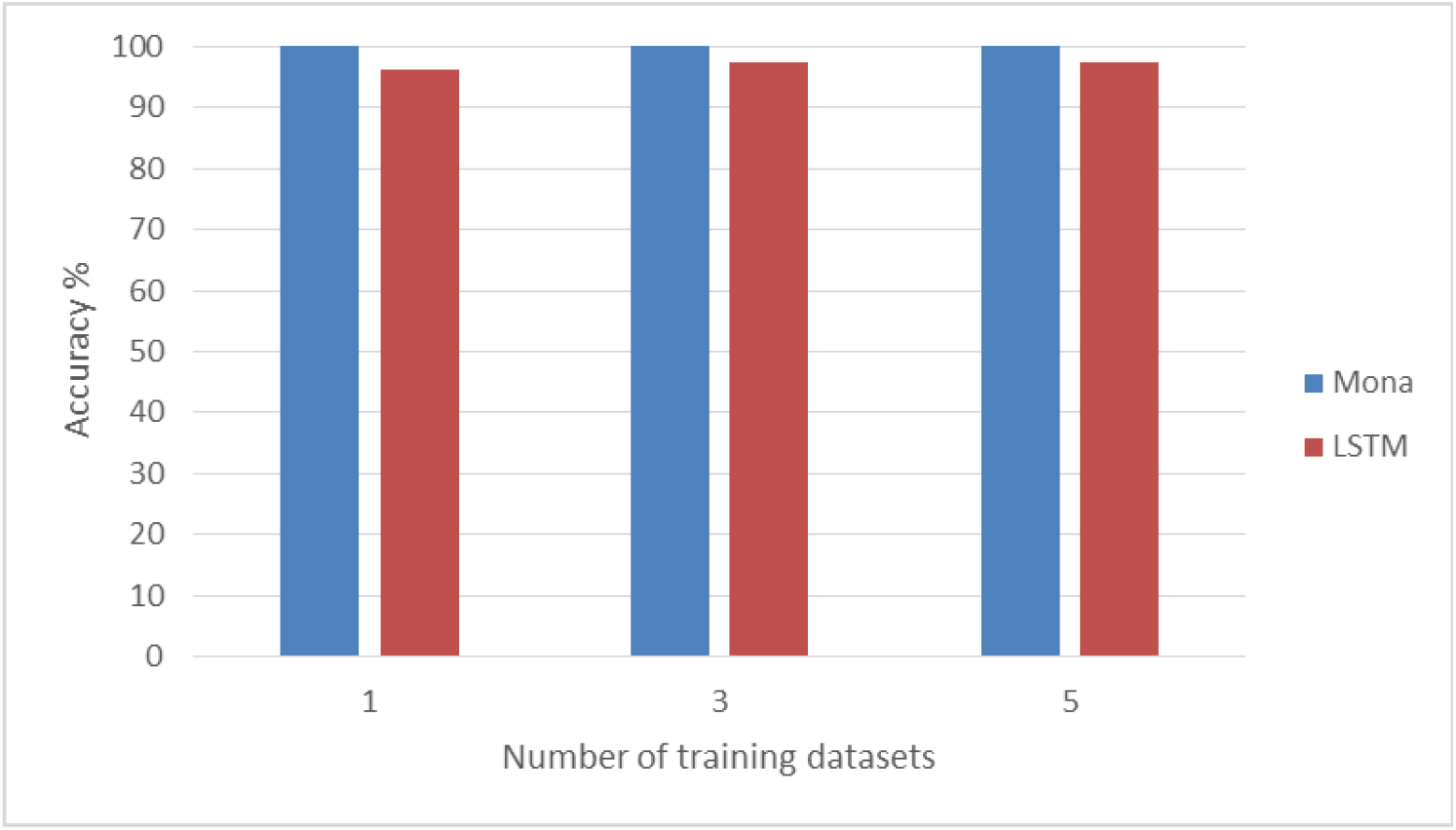
Male test performance with number of training datasets.

Each run is seeded with a different random number that controls the movements of the mice in the forest and how long eating a mouse serves to satisfy a bird’s hunger. The testing accuracy was calculated as the percent of correct responses out of 1000 steps. Measurements were based on the mean values of 20 trials. Very little variance in values was observed.

Both networks show excellent performance with only a single training dataset.

### Dynamic testing

Mona performance is measured as it interacts with the world. That is, responses output to the world cause changes in the world that are subsequently input to the bird. For example, if the bird turns to the left, the world reacts by altering the sensory state of the bird accordingly. With place motor neurons, Mona solves the task every time, but without place motors the male bird, which must perform complex navigation to fetch mice and stones, repeatedly becomes lost, causing the entire task to fail.

When the same “dynamic” interface is applied to the RNN network the male bird repeatedly becomes lost while fetching mice or stones. This means that even the few errors that the male makes are crucial, preventing successful completion. If the male cannot return with a mouse for the female, for example, the female cannot proceed with nest building.

### RNN epoch testing

Mona trains in a single epoch, a skill frequently seen in human learning (Lee et al., 2015). The RNN training is significantly affected by the number of epochs of training, especially for the male, as shown in Figure 10.

**Figure 10.**
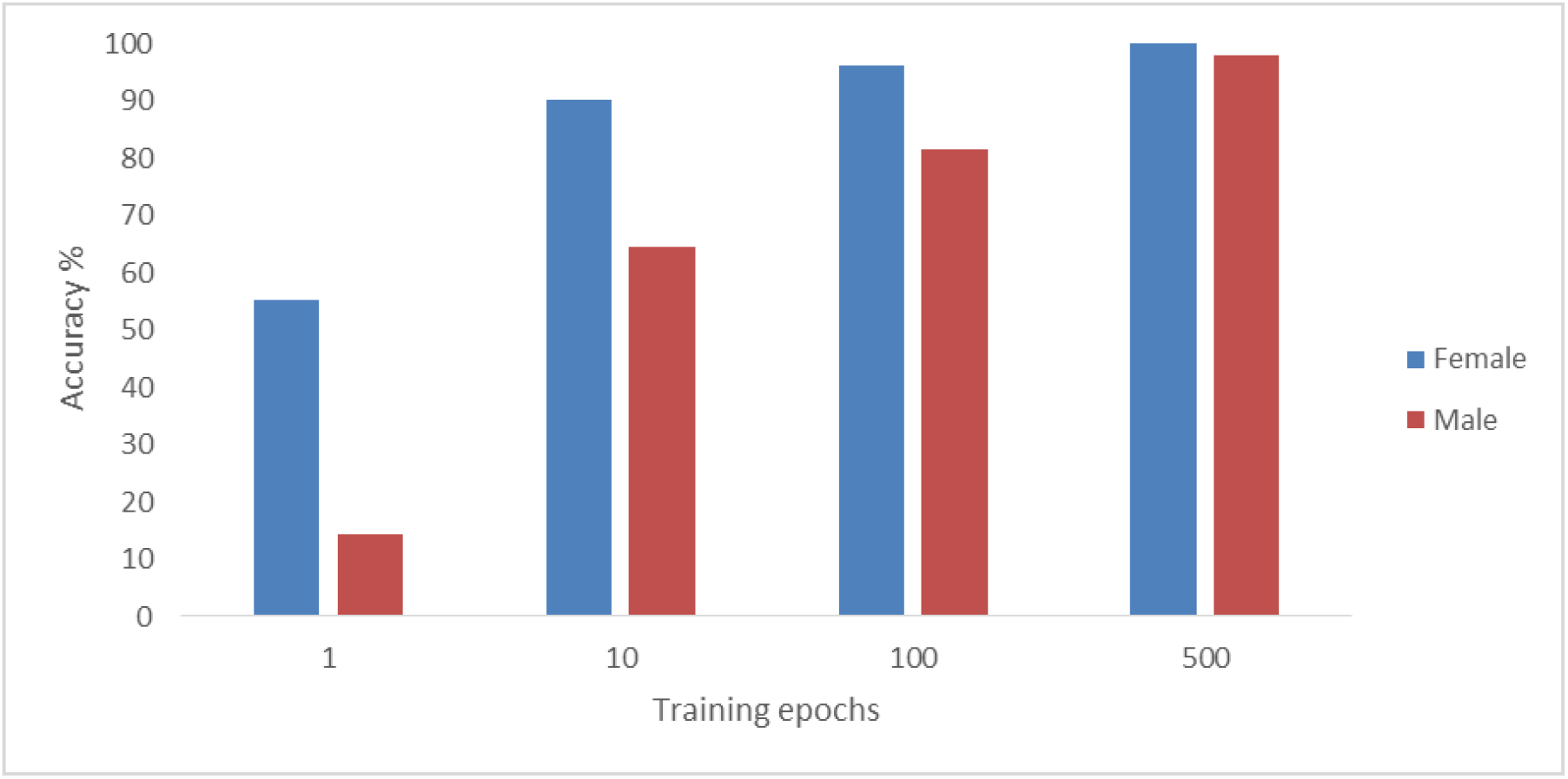
Training epochs testing

## Conclusion

Enhanced with place motor neurons, Mona is capable of solving the nest-building task every time. The RNN performs well with its typical usage, which is to predict upcoming responses, but fails as a navigator for the male bird, causing nest-building to be unsuccessful. How place neuron functionality might be incorporated into an RNN is an interesting topic.

Place motor neurons and goal-seeking causation learning are a powerful combination of capabilities for the nest-building task which demands both spatial and sequential learning. This animal learning task exposes shortcomings in a deep learning ANN that researchers interested in artificial general intelligence (AGI) should be aware of. I recommend that further research be conducted to (1) further simulate animal behaviors, and (2) adopt mechanisms from neurobiology, such as place cells, that allow machines to acquire animal-like capabilities. These I believe are essential to the achievement of AGI.

